# Unsupervised clustering analysis of SARS-Cov-2 population structure reveals six major subtypes at early stage across the world

**DOI:** 10.1101/2020.09.04.283358

**Authors:** Yawei Li, Qingyun Liu, Zexian Zeng, Yuan Luo

**Affiliations:** Department of Preventive Medicine, Northwestern, University, Feinberg School of Medicine, Chicago, IL 60611, USA; Department of Immunology and Infectious Diseases, Harvard T. H. Chan School of Public Health, Boston, MA 02115, USA; Department of Data Science, Dana Farber Cancer Institute, Harvard T. H. Chan School of Public Health, Boston, MA 02115, USA

**Keywords:** Deep learning clustering, population structure, evolution, SNP, SARS-CoV-2

## Abstract

Identifying the population structure of the newly emerged coronavirus SARS-CoV-2 has significant potential to inform public health management and diagnosis. As SARS-CoV-2 sequencing data accrued, grouping them into clusters is important for organizing the landscape of the population structure of the virus. Due to the limited prior information on the newly emerged coronavirus, we utilized four different clustering algorithms to group 16,873 SARS-CoV-2 strains, which automatically enables the identification of spatial structure for SARS-CoV-2. A total of six distinct genomic clusters were identified using mutation profiles as input features. Comparison of the clustering results reveals that the four algorithms produced highly consistent results, but the state-of-the-art unsupervised deep learning clustering algorithm performed best and produced the smallest intra-cluster pairwise genetic distances. The varied proportions of the six clusters within different continents revealed specific geographical distributions. In particular, our analysis found that Oceania was the only continent on which the strains were dispersively distributed into six clusters. In summary, this study provides a concrete framework for the use of clustering methods to study the global population structure of SARS-CoV-2. In addition, clustering methods can be used for future studies of variant population structures in specific regions of these fast-growing viruses.

## I. Introduction

The COVID-19 pandemic was caused by severe acute respiratory syndrome coronavirus 2 (SARS-CoV-2) [1, 2], and has spread throughout the world. In an effort to understand the molecular characteristics of the virus, viral genomes have been abundantly sequenced and presented at the Global Initiative on Sharing All Influenza Data (GISAID). As an emerging virus, it is important to understand the genetic diversity, evolutionary trajectory and possible routes of transmission of SARS-CoV-2 from its natural reservoir to humans. Most studies have looked into the aspects of real-world SARS-CoV-2 evolution and strain diversification through phylogenetic tree [3-7]. Phylogenetic tree is a graph that shows the evolutionary relationships among various biological entities based on their genetic closeness [8, 9]. The distances from one entity to the other entities indicate the degree of relationships. However, as population genomic datasets grow in size, simply using pairwise genetic distances cannot present an explicit structure of the total population in phylogenetic analysis. Grouping similar entities into the same cluster and identifying the number of main subtypes (clusters) makes it easier to understand the main characteristics of the population. Traditionally, using the distance matrix and the bifurcations between branches of leaves on the phylogenetic tree, entities can be grouped into clusters. However, when the number of entities becomes large, it is not easy to directly and accurately partition the clades in the phylogenetic tree.

To identify a better way to effectively group entities, clustering methods emerge as more productive and robust solutions. The objective of clustering is automatically minimizing intra-cluster distances and maximizing inter-cluster distances [10]. Accurate clustering helps to better understand the inner relationships between data and inform downstream analysis. Clustering methods have been widely used as a good supplemental tool in phylogenetic analysis, including phylogenetic tree construction [11-13], ancestral relationship identification [14], evolutionary rate estimation [15, 16], gene evolutionary mechanisms research [17] and population structure analysis [18].

Herein, to identify the population structure of the newly emerged coronavirus SARS-CoV-2, we took inspiration from recent state-of-the-art deep embedding clustering method [19] to group a total of 16,873 strains. Compared with traditional methods, this deep learning clustering algorithm showed significant improvements in terms of both Silhouette score, sum of squared errors (SSE) and Bayesian information criterion (BIC) [20]. The clustering results showed that there were six major clusters of SARS-CoV-2. In particular, we found that the proportions of six clusters in each continent showed a specific geographical distribution. In summary, this study provides a perspective of the SARS-CoV-2 population structural analysis, helping to investigate the evolution and spread of the virus across the human populations worldwide.

## II. METHODOLOGY

### A. SARS-CoV-2 sample collection

A set of African, Asian, European, North American, Oceanian and South American SARS-CoV-2 strains marked as “high coverage” were downloaded from GISAID. The “high coverage” was defined as strains with <1% Ns and <0.05% unique amino acid mutations (not seen in other sequences in databases) and no insertion/deletion unless verified by the submitter. In addition, all strains with a non-human host and all assemblies of total genome length less than 29,000 bps were removed from our analysis. Ultimately, our dataset consisted of 16,873 strains.

### B. Mutation calls and phylogenetic reconstruction

All downloaded genomes were mapped to the reference genome of SARS-CoV-2 (GenBank Accession Number: NC_045512.2) following Nextstrain pipeline [21]. Multiple sequence alignments and pairwise alignments were constructed using CLUSTALW 2.1 [22]. Considering many putatively artefactual mutations and the gaps in sequences are located at the beginning and end of the alignment, we masked the first 130 bps and last 50 bps in mutation calling following Nextstrain pipeline. We used substitutions as features to reconstruct the phylogenetic tree using FastTree 2 [23]. The phylogeny is rooted following Nextstrain pipeline using FigTree v1.4.4. The phylogenetic trees were visualized using the online tool Interactive Tree Of Life (iTOL v5) [24].

### C. Data analysis and visualization

All Figures and statistical analyses were generated by the ggplot2 library in R 3.6.1, the seaborn package in Python 3.7.6 and GraphPad Prism 8.0.2.

### D. Data clustering

Herein, we employed a published state-of-the-art deep learning unsupervised clustering algorithm to iteratively cluster the SARS-CoV-2 strains [19]. Each identified cluster was a subtype of SARS-CoV-2. We first used K-means clustering to initialize centroids for the clusters. To determine the number of clusters, we plotted the curves of the sum of squared errors (SSE) and Bayesian information criterion (BIC) [20] under different cluster numbers ranging from 2 to 20.

To update the cluster assignments, we implemented the Student’s t-distribution as a kernel to measure the distance from a strain (*h*_*i*_) to a cluster centroid (*u*_*j*_):

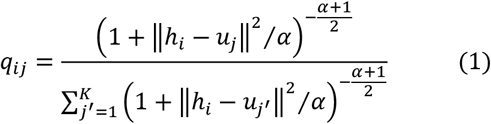

where the distance *q*_*ij*_ can be interpreted as the probability of assigning strain i to cluster j. The *α* is the degree of freedom of the Student’s t-distribution, and we let *α* = 1 in this study. Next, we defined an auxiliary target distribution P by raising each *q*_*ij*_ to the second power which upweights strains assigned with high confidence:

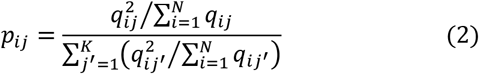

where the denominator is to normalize the loss contribution of each centroid to prevent large clusters from distorting the feature space. Finally, we defined the objective function using a Kullback-Leibler (KL) divergence loss:

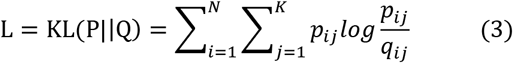

The parameters and cluster centroids were jointly optimized by minimizing L using Stochastic Gradient Descent (SGD) with momentum.

Besides the deep learning clustering algorithm, we also employed K-means clustering, hierarchical clustering and BIRCH (Balanced Iterative Reducing and Clustering using Hierarchies) for SARS-CoV-2 strain clustering. The three other models were implemented using the Python package sklearn with the KMeans function (K-means), AgglomerativeClustering function (hierarchical clustering) and Birch function (BIRCH), respectively.

### E. Data Availability

The publicly available SARS-CoV-2 datasets in this study are available at GISAID (https://www.gisaid.org). The reference SARS-CoV-2 is available at the NCBI GenBank (GenBank Accession Number: NC_045512.2, https://www.ncbi.nlm.nih.gov/nuccore/NC_045512.2).

## III. EXPERIMENTAL RESULTS

### A. Genetic analysis indicates high diversity and rapidly proliferating of SARS-CoV-2

We obtained a total of 16,873 (98 from Africa, 1324 from Asia, 9527 from Europe, 4765 from North America, 1040 from Oceania and 119 from South America) earliest SARS-CoV-2 whole-genome sequencing data from GISAID, aligned the sequences, and identified the genetic variants. A total of 7,970 substitutions were identified, including 4,908 non-synonymous mutations, 2,748 synonymous mutations and 314 intronic mutations. The average mutation count per genome was 6.99 (Fig. 1A). The frequency spectrum of substitutions illustrated that more than half (54.05%) of the mutations were singletons and 15.35% were doubletons. The proportion of the mutations below 0.01 was 99.28% (Fig. 1B). The high percentage of these low-frequency mutations suggested that SARS-CoV-2 occurred recently and displayed a rapidly proliferating pattern [25]. In addition, there were 8,706 unique strains across the 16,873 strains (Fig. 1C), and most unique strains (7,078) were singletons, yielding high diversity of the virus. In particular, Simpson’s diversity index of the strains was 0.8222, indicating that two random strains would have a high probability of being genetically different. The frequency spectrum of substitutions and Simpson’s diversity index indicated high genetic diversity of SARS-CoV-2.

**Fig. 1.**
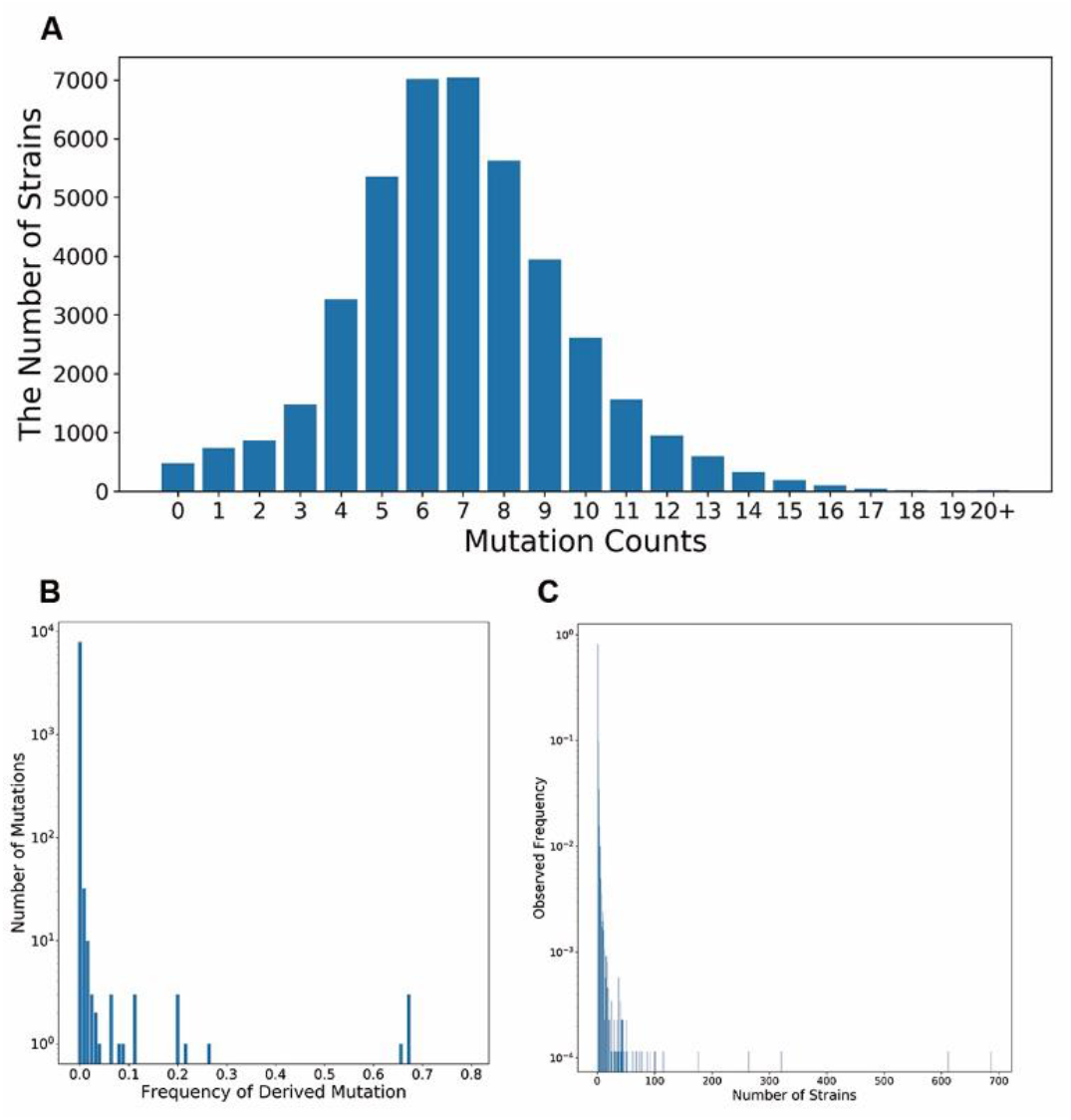
The genetic information of the 16,873 SARS-CoV-2 strains. (A) The distribution of the mutation counts of SARS-CoV-2. (B) Frequency spectra of SARS-CoV-2. The mutation frequency of derived mutations of 16,873 SARS-CoV-2 stains is depicted on the X axis, and the number of mutations in corresponding strains occurred is displayed on the Y axis. A log-10 scale is used for the Y axis of the graph, and the Y axis ranges from 1 to 10,000. (C) Normalized allele frequency of SARS-CoV-2. There are 8,706 unique genomes across the 16,873 strains. The X axis is the number of strains for each unique genome and the Y axis is the proportion of the unique genomes. A log-10 scale is used for the Y axis of the graph, and the Y axis ranges from 0.0001 to 1

### B. Clustering of SARS-CoV-2 reveals six major clusters

To clarify the main population structure of the virus, grouping these strains into clusters is necessary, as these clusters displayed the major types of the virus. However, the genetic analysis of SARS-CoV-2 showed that there were 8,706 unique strains across the 16,873 strains (Fig. 1C), it is not easy to directly and accurately partition the strains. For this reason, we applied clustering techniques to measure similarities between these strains and effectively group them.

Because SARS-CoV-2 exhibits a limited number of SNPs per virus strain and little ongoing horizontal gene exchange, making SNPs ideal clustering input features. We first used the aggregated SNP matrix to cluster samples using an unsupervised deep learning clustering algorithm published by Xie et al [19] (see methodology). The unsupervised deep learning clustering algorithm requires one to pre-specify the number of clusters (*K*), but we have little prior knowledge about the number of subtypes formed by the heterogeneous SARS-CoV-2 genome. To determine the number of clusters, we plotted the curves of the SSE and BIC under different cluster numbers ranging from 2 to 20 (Fig. 2). We used the elbow method and chose the elbow of the curve as the number of clusters [26]. This approach resulted in *K*=6 for both the SSE and BIC curves.

**Fig. 2.**
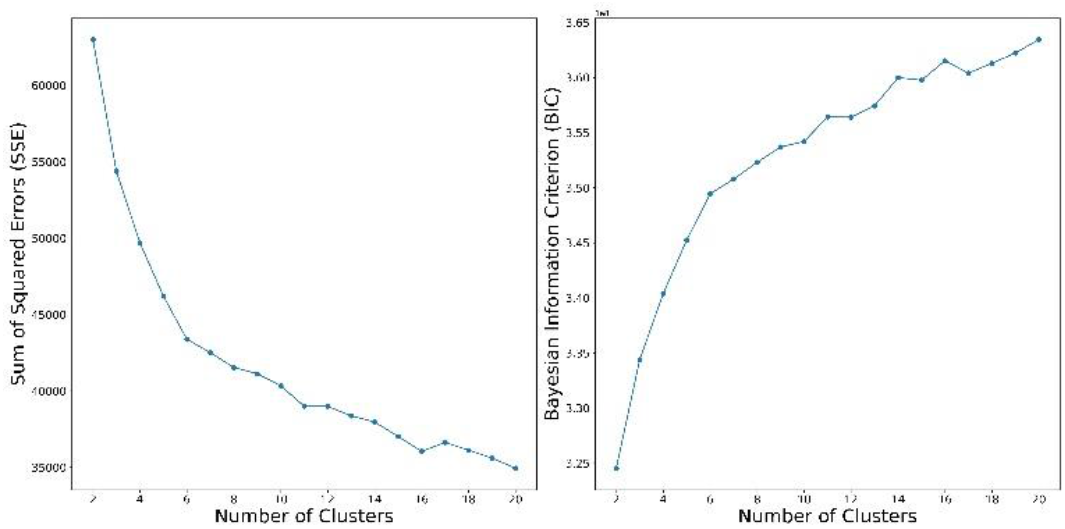
Evaluation of the number of clusters. The evolution of the sum of squared errors (SSE; left) and Bayesian information criterion (BIC; right) for the number of clusters in the deep learning clustering runs. We used the elbow method and chose the elbow of the curve as the number of clusters. The elbow method indicated that the number of clusters is six.

To evaluate the performance of the algorithm, we also employed K-means clustering [27], hierarchical clustering and BIRCH clustering [28, 29] for comparison. The objective of clustering is minimizing intra-cluster distances and maximizing inter-cluster distances. To this end, we did five repetitions for each of the four clustering algorithms and selected the one that achieved the best performance (lowest average intra-cluster pairwise genetic distances). The average intra-cluster pairwise genetic distances in the deep learning clustering algorithm (4.892) was significantly lower than that in K-means (4.896, P-value < 0.001, Wilcoxon rank-sum test), hierarchical clustering (5.062, P-value < 0.001, Wilcoxon rank-sum test) and BIRCH (4.985, P-value < 0.001, Wilcoxon rank-sum test). We compared the Silhouette score (Fig. 3A), SSE (Fig. 3B) and BIC (Fig. 3C) of the four algorithms. The deep learning clustering obtained the highest Silhouette score and BIC, and the lowest SSE, indicating that the clustering results of deep learning clustering are better than the other algorithms. In contrast, BIRCH performed the worst of the four algorithms. We aligned the partitions of the six clusters against the phylogenetic tree for the three best methods (Fig. 3D). The clustering results indicated that the partitions from the three algorithms were similar. The differences between the hierarchical clustering results and the two other clustering results were mainly at the boundary of the clusters. Of the three methods, strains grouped by deep learning clustering and K-means were more compact in the phylogenetic tree than those by hierarchical clustering. For example, the strains in both deep learning clustering cluster D and K-means cluster D were split into two clusters using hierarchical clustering. However, such a split was not supported by the phylogenetic tree (Fig. 3D).

**Fig. 3.**
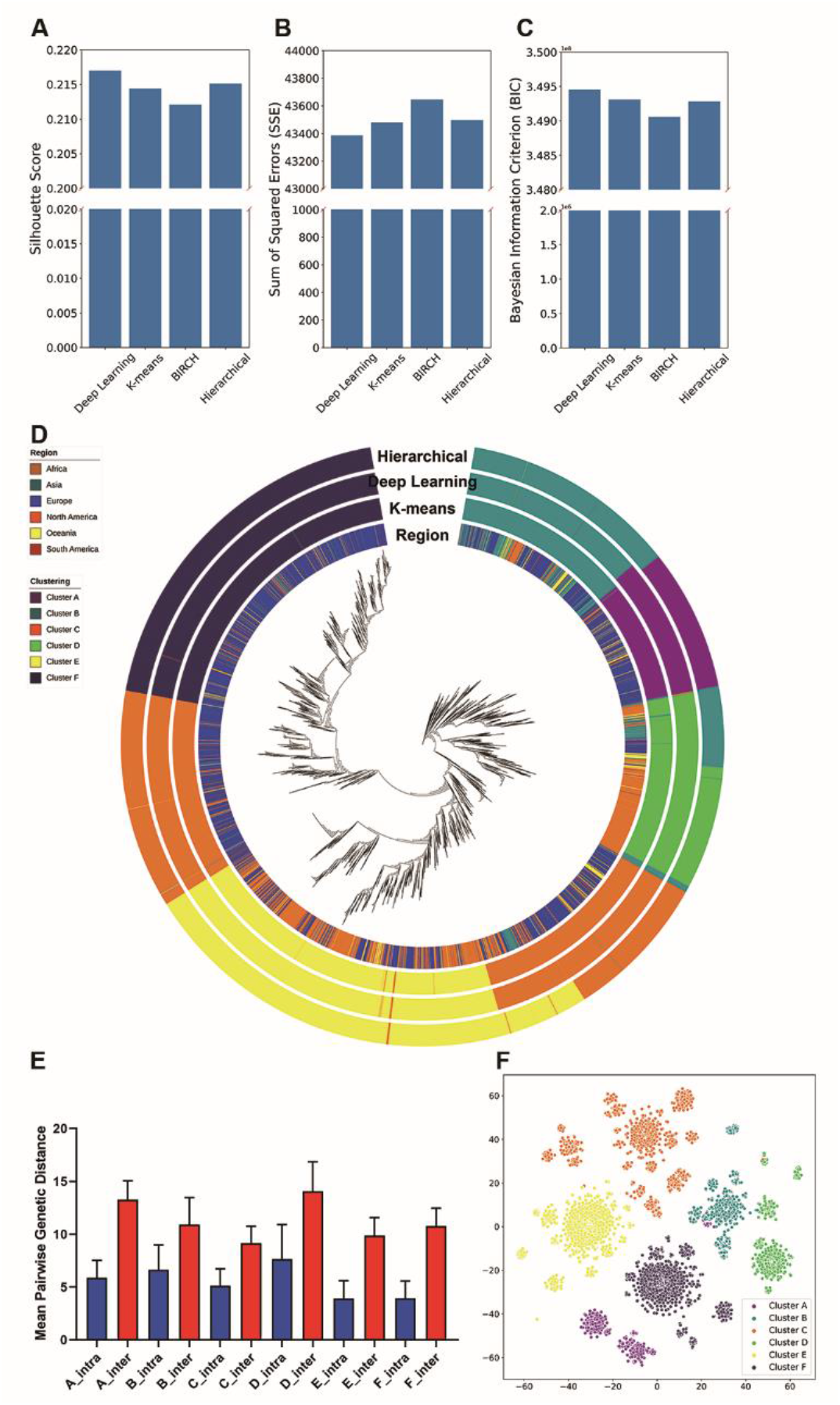
Clustering of SARS-CoV-2. (A, B and C) The Silhouette score (A), Sum of Squared Errors (SSE; B) and Bayesian Information Criterion (BIC; C) for the four selected algorithms (X axis). (D) Phylogenetic tree of 16,873 SARS-CoV-2 strains. Four colored panels outside the phylogenetic tree are used to identify auxiliary information for each virus strain. The inner panel represents the distribution of the continents. The outer three panels represent the partitions of the six clusters across the three best performance clustering algorithms (deep learning, K-means and Hierarchical) in the tree. (E) Mean pairwise genetic distances for intra-clustered and inter-clustered genetic distances. The blue bars represent mean pairwise genetic distances between pairs of isolates within the clusters, and the red bars represent mean pairwise genetic distances between pairs of isolates outside the clusters. The error bar represents the standard deviation. The mean distance between pairs of strains for intra-clusters was significantly lower (P-value < 0.001, Wilcoxon rank-sum test) than that of inter-clusters. (F) The t-SNE plot of the deep learning clustering results. Each dot represents one strain and each color represents the corresponding cluster.

In the meantime, we used complementary approaches to validate the deep learning clustering results. First, we compared the pairwise genetic distances between intra-cluster and inter-cluster. In all six clusters, the average number of intra-cluster genetic distances was significantly lower (P-value < 0.001, Wilcoxon rank-sum test, Fig. 3E) than inter-cluster genetic distances. Next, we applied T-distributed Stochastic Neighbor Embedding (t-SNE) to visualize the deep learning clustering results. In the t-SNE plot, the strains were adequately isolated between clusters (Fig. 3F).

### C. The varied proportions of the clusters in different continents

Mapping the proportions of strains from each continent showed that the clusters differed in their geographical distributions (Table 1). Of the six clusters, cluster C spread globally. By contrast, cluster A and cluster F occurred at high frequencies in specific regions. 81.92% of the strains in cluster A and 85.73% of the strains in cluster F were from Europe. The geographical spread of each of the three remaining clusters was intermediate. Cluster E occurred at higher frequencies in North America and Europe, lower frequencies in Asia and Oceania. Cluster D occurred at higher frequencies in North America, and lower frequencies in Asia, Europe and Oceania. The strains in cluster B were mainly in Asia and Europe and partially in North America and Oceania.

**Table 1.**
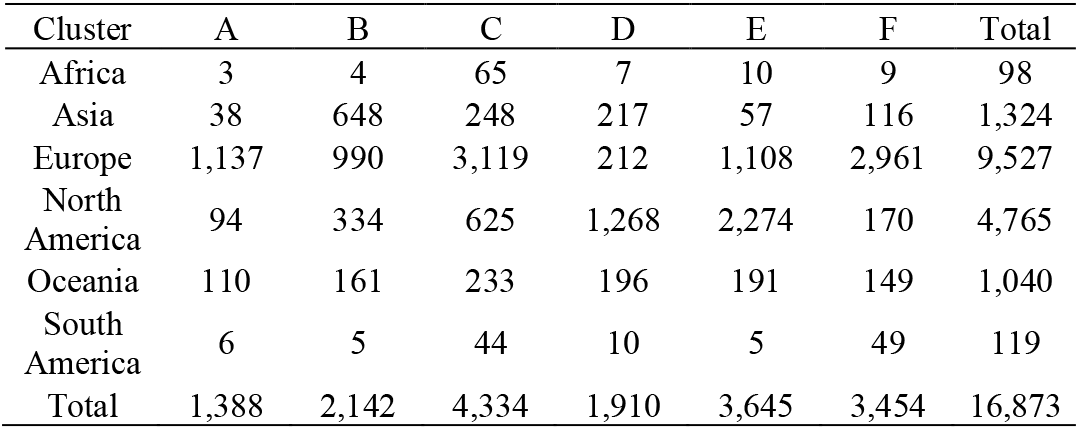
Geographic distribution of six continents for each cluster.

| Cluster | A | B | C | D | E | F | Total |
| --- | --- | --- | --- | --- | --- | --- | --- |
| Africa | 3 | 4 | 65 | 7 | 10 | 9 | 98 |
| Asia | 38 | 648 | 248 | 217 | 57 | 116 | 1,324 |
| Europe | 1,137 | 990 | 3,119 | 212 | 1,108 | 2,961 | 9,527 |
| North America | 94 | 334 | 625 | 1,268 | 2,274 | 170 | 4,765 |
| Oceania | 110 | 161 | 233 | 196 | 191 | 149 | 1,040 |
| South America | 6 | 5 | 44 | 10 | 5 | 49 | 119 |
| Total | 1,388 | 2,142 | 4,334 | 1,910 | 3,645 | 3,454 | 16,873 |

However, due to the sampling bias of the SARS-CoV-2, 85% of the strains were collected from Europe and North America, making the proportion of the continents in each cluster not informative. Therefore, we evaluated the proportion of the clusters on each continent. In most continents, the distributions of the strains were concentrated in one or two clusters, including Asia (49% in cluster B), Africa (66% in cluster C), South America (78% in cluster C and F), North America (74% in cluster D and E) and Europe (64% in cluster C and F). Among the six continents, Oceania was the only continent that was uniformly separated into the six clusters, indicating strains in Oceania were more diverse than in the other continents.

## IV. Conclusion

Understanding the population structure of SARS-CoV-2 is important in evaluating future risks of novel infections. To precisely analyze their population structure, we used clustering methods in phylogenetic analysis to group a total of 16,873 publicly available SARS-CoV-2 strains. To improve the accuracy, we use a state-of-the-art deep learning clustering algorithm, which has been demonstrated to exhibit better performance than three traditional clustering algorithms: K-means clustering, hierarchical clustering and BIRCH.

Our clustering results indicated six major clusters of SARS-CoV-2. The mutation profile characterizing clusters of the viral sequences displayed specific geographical distributions. Most continents were mainly concentrated in one or two clusters, but we also found that in Oceania, the strains were dispersively distributed into six clusters.

It is noteworthy that our study is limited due to the sampling bias of SARS-CoV-2, with more than 60% of the strains being from the United Kingdom and the USA. In contrast, the overall proportion of strains from Africa and South America is less than 2%. Sampling biases can lead to biased parameter estimation and affect the clustering results we observed. To address this issue, another clustering can be used for more further analyses.

Despite the limited number of SARS-CoV-2 genome sequences, our analysis of population genetics is informative. Our discovery of high genetic diversity in SARS-CoV-2 is consistent with an earlier study [30]. The topology and the divergence of the clusters in the phylogenetic tree illustrate a relatively recent common ancestor, similar to the fact that the emergence and the spread of the virus was highly concentrated in a short time [1, 31-33]. Our work, as well as previous studies [3, 34, 35] that use clustering techniques to study the population structure of the SARS-CoV-2 virus, has proved to be a valuable supplemental tool in phylogenetic analyses. For future work, we plan to further apply soft clustering techniques to better account mixtures in clusters, the efficacy of which has been showcased by previous studies in multiple fields [36-42]. In addition, clustering ideas can be used for further study of variant population structures in specific regions of these fast-growing viruses.

## Acknowledgment

We thank Xin Wu for the comments and suggestions during the preparation of the manuscript. This study is supported in part by NIH grant 1R01LM013337 and 5UL1TR001422.

